# Co-option of ancestral signal elements in the evolution of a cooperative parental behavior

**DOI:** 10.1101/2022.10.10.511585

**Authors:** Jeanette B. Moss, James P. Tumulty, Eva K. Fischer

**Affiliations:** Department of Evolution, Ecology, and Behavior, University of Illinois, Urbana-Champaign, Urbana, IL 61801, U.S.A.; Department of Biology, College of William and Mary, Williamsburg, VA, 23185, U.S.A.

**Author notes:** Eva K. Fischer, **Email:**. **Author Contributions:** J.B.M., J.P.T., and E.K.F. conceived of and designed the study. J.B.M. collected the data. J.B.M. and J.P.T. performed the analyses. J.B.M., J.P.T., and E.K.F. wrote the paper.

**Keywords:** Animal communication, anuran, parental care, cooperation, social context

## Abstract

The emergence of complex social interactions is predicted to be an important selective force in the diversification of communication systems. Parental care presents a key social context in which to study the evolution of novel signals, as care often requires communication and behavioral coordination between parents and is an evolutionary stepping-stone towards increasingly complex social systems. Anuran amphibians (frogs and toads) are a classic model of acoustic communication and the vocal repertoires of many species have been characterized in the contexts of advertisement, courtship, and aggression; yet quantitative descriptions of calls elicited in the context of parental care are lacking. The biparental poison frog, *Ranitomeya imitator*, exhibits a remarkable parenting behavior in which females, cued by the calls of their male partners, feed tadpoles unfertilized eggs. Here, we characterized and compared calls across three social contexts, for the first time including a parental care context. We found that egg feeding calls share some properties with both advertisement and courtship calls but also had unique properties. Multivariate analysis revealed high classification success for advertisement and courtship calls but misclassified nearly half of egg feeding calls as either advertisement or courtship calls, suggesting additional signal modalities play a role in parental communication. Egg feeding and courtship calls both contained less identity information than advertisement calls, as expected for signals used in close-range communication where uncertainty about identity is low. Taken together, egg feeding calls likely borrowed and recombined elements of both ancestral call types to solicit a novel, context-dependent parenting response.

**Significance Statement:** Parental care has evolved independently in every major animal lineage and represents a major step in the evolution of complex sociality. Communication systems may need to increase in complexity. To explore these ideas, we characterized calls associated with trophic egg feeding, a unique cooperative parental behavior in the biparental mimic poison frog and compared them to calls associated with mate attraction (advertisement and courtship calls). Our analysis revealed some distinct, but many shared properties of signals elicited during egg feeding, suggesting that signals deployed in a novel social context evolve via modification and recombination of existing signals. These findings deepen our understanding of the relationship between complexity of social and communication systems.

## Introduction

Complexity of communication systems is predicted to co-evolve with complexity of social systems (1, 2). Transitions in social organization introduce novel contexts for interaction between individuals, which in turn require more specialized communication systems. While traditionally examined through the lens of group size (3–5), there is increasing interest in understanding how these co-evolutionary dynamics play out across multiple indices of social complexity (6–8). Parental care is a major component of a species’ social system (9) and represents a key step in the evolution of complex sociality (10). When care responsibilities are shared between multiple caregivers, ensuring offspring survival requires coordinated behaviors that rely on effective communication. One solution is to expand the size of the signal repertoire by adding novel, context-specific communication features (7, 11), as has been shown in the case of avian cooperative breeding (8). Alternatively, existing signals may be co-opted to function in the new context via context-dependent regulation of responses (12, 13). Quantitative descriptions of variation in signal components provide an important first step for generating hypotheses about signal function and evolution.

One of the most influential models for studying animal communication is anuran amphibian vocal communication and auditory processing. Across anurans, vocal repertoires of differing sizes and compositions have been meticulously characterized along with the diversity of behavioral and neural responses they elicit reviewed in 14–16). This rich literature illustrates how greater complexity of communication systems may evolve through the specialization of signal components for a given behavioral context, evolution of receiver bias, or both. In a classic example, the common coquí frog (*Eleutherodactylus coquí*), advertisement call is composed of two syllables – ‘co’ and ‘qui’ – which are differentially specialized for aggressive versus mate attraction contexts (17). In this case, regulation of responses is further facilitated by the evolution of sex-specific auditory sensitivities (17). Conversely, modulation of the túngara frog’s (*Physalaemus pustulosus*)characteristic ‘whine-chuck’ call has similar motivating effects on potential mates and rivals, but with clearly distinct social and behavioral outcomes (18).

Beyond serving as models for the study of animal communication, anurans also exhibit remarkable diversity of parental care (19). The mimic poison frog, *Ranitomeya imitator*, is a species characterized by monogamy, pair bonding, and biparental care of eggs and tadpoles (20). Terrestrial eggs are cared for by both parents and tadpoles are transported “piggy-back” to pools of water upon hatching. Remarkably, parents visit and care for their developing tadpoles until metamorphosis. Specifically, females feed tadpoles with unfertilized ‘trophic’ eggs, a fascinating behavior coordinated by males calling to females and leading them to hungry, begging tadpoles. Egg feeding has evolved multiple times in association with biparental care (20) and functions crucially in offspring development (21), suggesting that selection on signalers and receivers during these interactions is strong. It has been hypothesized that egg feeding behaviors are evolutionarily derived from courtship behaviors (21, 23, 24); however, the signals that coordinate and elicit egg feeding have yet to be quantitatively described for any species, including *R. imitator*.

Here, we characterize and compare calls of *R. imitator* across three social contexts to test whether the emergence of coordinated parental care within this lineage has promoted the incorporation of novel signals for communication. Given that several call types in frogs are known to serve dual functions, we predicted that egg feeding calls were likely co-opted and modified from calls adapted for the more ancestral functions of advertisement and courtship (described in 29 and 30).

## Results

### Calls vary across social contexts

To quantify and compare advertisement, courtship, and egg feeding signals, we recorded representative calls of each type for captive male *Ranitomeya imitator* across the breeding cycle. Briefly, advertisement and courtship calls – both deployed for attracting females but over different distances (i.e., long vs. short range, respectively; 6, 7, 33) were distinguished based on published descriptions of *R. imitator* vocalizations (25, 26), whereas egg feeding calls were identified using video surveillance of parenting frogs (e.g., Movie S1). Visual inspection of waveforms and spectrograms of putative call types revealed distinct characteristics (Fig. 1).

**Figure 1.**
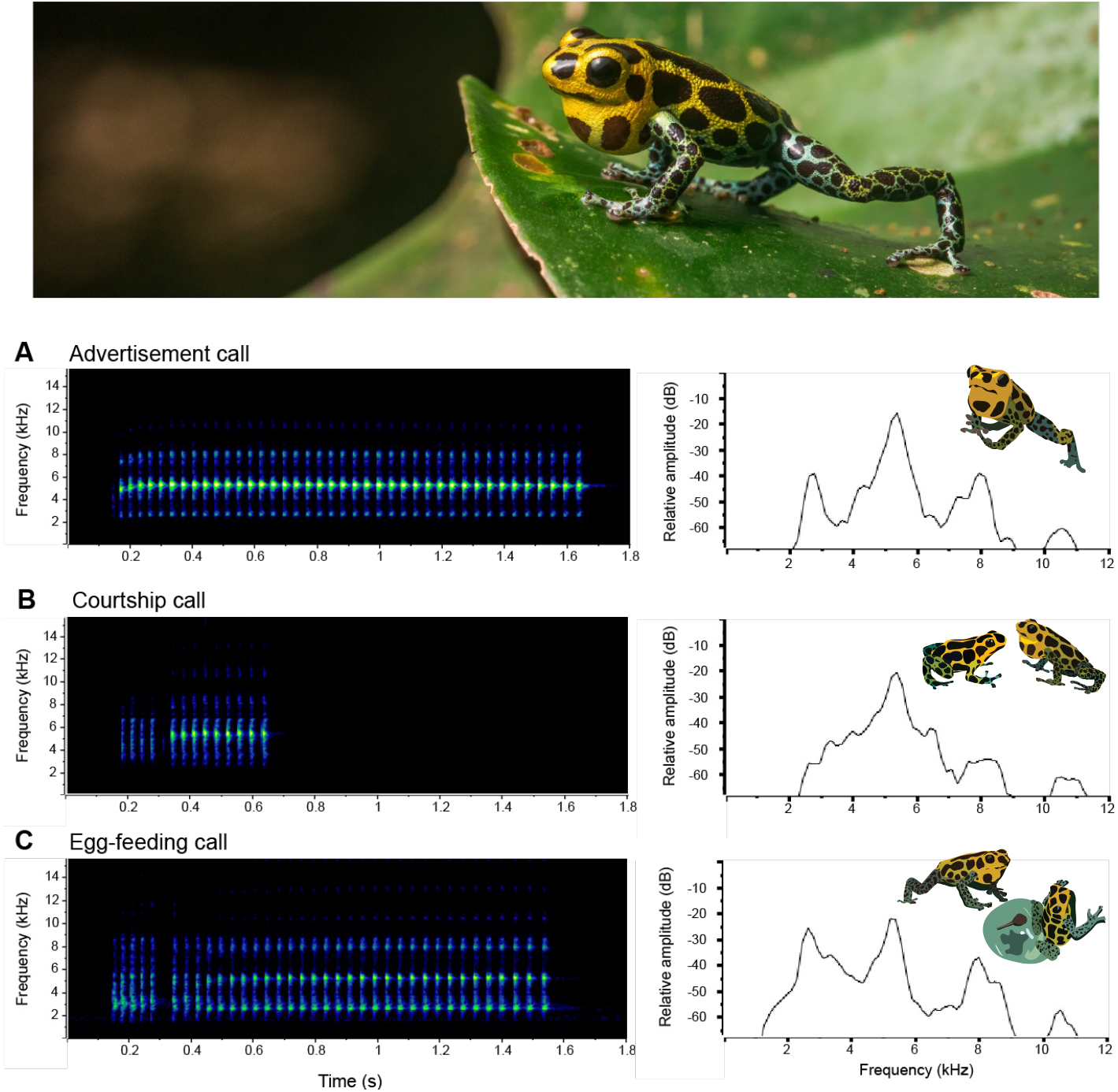
Vocalizations have unique spectral profiles across social contexts. Spectrograms and associated power spectra of a representative (A) advertisement call; (B) courtship call; and (C) egg feeding call of *Ranitomeya imitator* (photo, top). Calls were visualized in Raven Pro v. 1.6.1 (Cornell Lab of Ornithology, 2022) using the Hann window, 256 pt frequency resolution. The courtship and egg feeding call power spectra show the two forms of increased frequency bandwidth: broader dominant frequency peak (B) or a fundamental frequency that is at a higher relative amplitude (C). Photo by Anton Sorokin.

Because signals are composed of multiple components, which may differ in salience to different receivers and be under different forms of selection, we measured nine acoustic properties to quantify call variation (Table 1; Fig. S1). All properties except for inter-call interval and pulse rate showed significant variation depending on the social context in which the call was measured (ANOVA *p* < 0.05). In terms of temporal properties, courtship calls were significantly shorter than advertisement (t = 6.59, *p* < 0.001) or egg feeding calls (t = 4.56, *p* < 0.001) and contained fewer pulses (advertisement-courtship: t = 7.24, *p* < 0.001; courtship-egg feeding: t = 4.98, *p* < 0.001). Pulse durations were longest in advertisement calls (advertisement-courtship: t = 3.06, *p* = 0.003; advertisement-egg feeding: t = 4.65, *p* < 0.001) and shortest in egg feeding calls (egg feeding-courtship: t = 2.52, *p* = 0.013; Fig. 2A), while pulse intervals were shortest in advertisement calls (advertisement-egg feeding: t = 4.22, *p* < 0.001; advertisement-courtship: t = 2.72, *p* = 0.007;) and longest in egg feeding calls (courtship-egg feeding: t = 2.33, *p* = 0.021) (Fig. 2B). In terms of spectral properties, egg feeding calls had significantly lower dominant frequencies than either advertisement (t = 3.67, *p* < 0.0001) or courtship calls (t = 2.88, *p* = 0.004; Fig. 2C), and advertisement calls had significantly narrower call bandwidths than courtship (t = 6.93, *p* < 0.001) or egg feeding calls (t = 5.77, *p* < 0.001; Fig. 2D).

**Figure 2.**
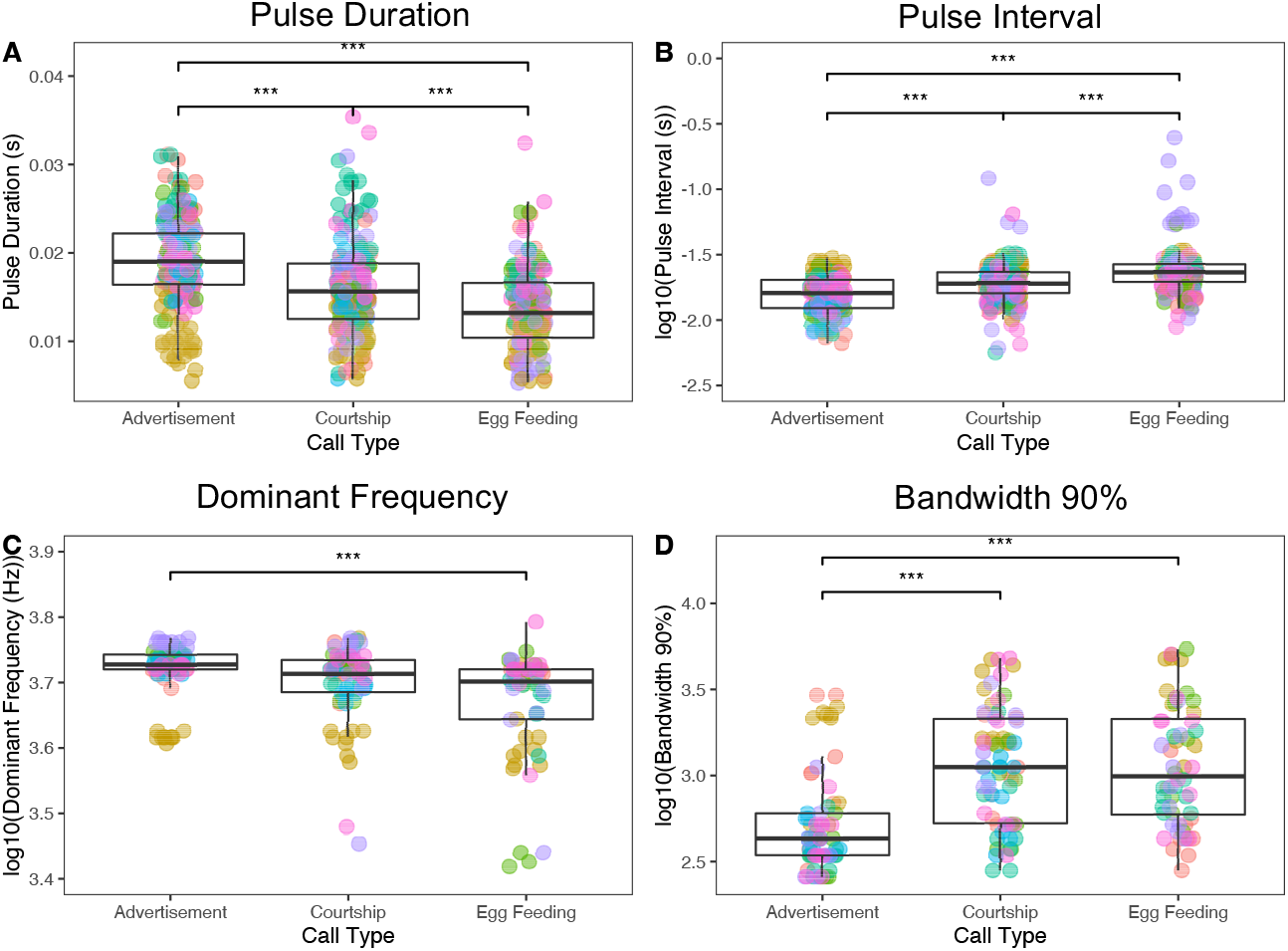
Acoustic properties of *Ranitomeya imitator* vocalizations vary between advertisement, courtship, and egg feeding calls. Plots depict distributions of four acoustic properties by call type: (A) Pulse duration; (B) log10(Pulse interval); (C) log10(Dominant frequency); and (D) log10*Bandwidth 90%). Color of dots corresponds to seven unique males in the sample. Significance levels are shown for Tukey-HSD post-hoc contrasts. Pairwise contrasts without significant differences are not shown.

**Table 1:** Summary of acoustic properties of 70 advertisement calls, 70 courtship calls, and 60 egg feeding calls across seven males. Pulse duration, interval, and rate were measured three times for each call, and summary statistics are thus calculated based on 210 or 180 measurements, respectively. Results of pairwise contrasts between call types are also reported. Tukey test statistics (t) and adjusted *p*-values are reported for each contrast and significant test-statistics are bolded.

| Acoustic Property | Advertisement | Courtship | Egg feeding | Advertisement-Courtship |  | Advertisement-Egg feeding |  | Courtship-Egg feeding |  |
| --- | --- | --- | --- | --- | --- | --- | --- | --- | --- |
| | Mean (± SD) | Mean (± SD) | Mean (± SD) | t | *p* | t | *p* | t | *p* |
| *Temporal Properties* |  |  |  |  |  |  |  |  |  |
| Call Duration (s) | 1.14 ± 0.23<br>(0.66–1.73) | 0.75 ± 0.29<br>(0.14–1.92) | 1.04 ± 0.35<br>(0.48–1.97) | **6.589** | **<0.001** | 1.072 | 0.285 | **-4.557** | **<0.001** |
| Call Interval (s) | 10.53 ± 7.93<br>(2.53–40.89) | 8.15 ± 5.42<br>(0.41–34.96) | 7.40 ± 5.27<br>(1.46–28.94) | 1.568 | 0.119 | 1.424 | 0.156 | 0.198 | 0.843 |
| No. of Pulses | 31.33 ± 6.02<br>(18–46) | 20.93 ± 7.28<br>(5–39) | 27.08 ± 9.22<br>(7–48) | **7.235** | **<0.001** | 1.164 | 0.246 | **-4.979** | **<0.001** |
| Pulse Duration (s) | 0.019 ± 0.005<br>(0.005–0.031) | 0.016 ± 0.005<br>(0.006–0.035) | 0.014 ± 0.005<br>(0.005–0.032) | **3.064** | **0.003** | **4.645** | **<0.001** | **2.519** | **0.013** |
| Pulse Interval (s) | 0.016 ± 0.005<br>(0.007–0.030) | 0.02 ± 0.01<br>(0.006–0.120) | 0.027 ± 0.023<br>(0.009–0.247) | **-2.722** | **0.007** | **-4.222** | **<0.001** | **-2.330** | **0.021** |
| Pulse Rate (notes/s) | 28.55 ± 2.16<br>(19.8–37.5) | 28.16 ± 3.67<br>(7.6–37.9) | 26.73 ± 5.02<br>(3.9–38.8) | 0.990 | 0.324 | 1.907 | 0.058 | 1.267 | 0.207 |
| *Spectral Properties* |  |  |  |  |  |  |  |  |  |
| Dominant Frequency (Hz) | 5249.1 ± 496.2<br>(4048.2–5857) | 5040.3 ± 613<br>(2842.4–5857) | 4783.3 ± 779.1<br>(2625–6201.6) | 1.403 | 0.162 | **3.676** | **<0.001** | **2.879** | **0.004** |
| Frequency of 2<sup>nd</sup> Harmonic (Hz) | 5220.8 ± 601.1<br>(4048.2–5857) | 5122.1 ± 495.2<br>(3789.9–5857) | 4954.8 ± 542.1<br>(3617.6–6201.6) | 1.520 | 0.130 | **3.946** | **<0.001** | **3.080** | **0.002** |
| Bandwidth 90% (Hz) | 678.8 ± 675.8<br>(258.4–2928.5) | 1544.1 ± 1193.4<br>(281.3–4823.4) | 1662.7 ± 1419.7<br>(281.3–5426.4) | **-6.932** | **<0.001** | **-5.767** | **<0.001** | -0.413 | 0.680 |

### Individual identity information varies across call types

We next examined the degree to which acoustic properties of each call type convey information about signaler identity. We found that putative call types and acoustic properties within those call types varied in their individual distinctiveness (Table 2). A key criterion for conveying information about signaler identity is that signal properties exhibit high levels of variation among individuals (CV_a_) but low levels of variation within individuals (CV_w_; 36). For calls classified as advertisement calls, most acoustic properties showed greater variation among individuals than within individuals (CV_a_ / CV_w_, or potential for individual coding (PIC) > 1). The greatest individual distinctiveness was found in spectral properties, in particular dominant frequency (PIC = 5.50). Courtship and egg feeding calls showed less individual distinctiveness overall, with most acoustic properties showing as much or more variation within individuals as among individuals, and dominant frequency being a less reliable indicator of identity (courtship PIC = 0.63; egg feeding PIC = 0.82). When considering all acoustic properties together, courtship calls contained the least identity information (*Hs* = 1.48) of the three types (advertisement *H_s_* = 3.82; egg feeding calls *H_s_* = 2.81).

**Table 2:** Individual distinctiveness of nine acoustic properties of 70 advertisement calls, 68 courtship calls, and 56 egg feeding calls across seven males. Potential for individual coding (PIC) is calculated as the ratio of the variation among individuals to the mean variation within individuals. Values greater than one indicate that a call property varies more among individuals than within individuals (shown in bold).

| Acoustic Property | Advertisement |  |  | Courtship |  |  | Egg feeding |  |  |
| --- | --- | --- | --- | --- | --- | --- | --- | --- | --- |
|  | CV<sub>a</sub> | Mean CV<sub>w</sub> | PIC | CV<sub>a</sub> | Mean CV<sub>w</sub> | PIC | CV<sub>a</sub> | Mean CV<sub>w</sub> | PIC |
| ***Temporal Properties*** |  |  |  |  |  |  |  |  |  |
| Call Duration (s) | 11.76 | 16.74 | 0.70 | 13.06 | 34.36 | 0.38 | 20.67 | 26.24 | 0.79 |
| Interval to next call (s) | 29.56 | 65.90 | 0.45 | 36.30 | 50.91 | 0.71 | 44.70 | 57.25 | 0.78 |
| No. of Pulses | **15.39** | **13.43** | **1.15** | 18.77 | 30.32 | 0.62 | **26.43** | **23.62** | **1.12** |
| Pulse Duration (s) | **20.29** | **19.48** | **1.04** | 18.88 | 26.50 | 0.71 | 20.22 | 27.81 | 0.73 |
| Pulse Interval (s) | 22.21 | 23.45 | 0.95 | 8.43 | 38.56 | 0.22 | **40.16** | **35.16** | **1.14** |
| Pulse Rate (notes/s) | 5.88 | 7.59 | 0.77 | 5.84 | 11.02 | 0.53 | 9.34 | 13.95 | 0.67 |
| ***Spectral Properties*** |  |  |  |  |  |  |  |  |  |
| Dominant Frequency (Hz) | **9.94** | **1.81** | **5.50** | 6.21 | 10.09 | 0.63 | 9.94 | 12.10 | 0.82 |
| Frequency of 2<sup>nd</sup> Harmonic (Hz) | **11.35** | **3.81** | **2.98** | **7.37** | **6.49** | **1.14** | **9.6** | **6.58** | **1.46** |
| Bandwidth 90% (Hz) | **70.83** | **47.50** | **1.49** | 48.57 | 62.18 | 0.78 | 58.74 | 62.49 | 0.94 |

### High call type classification success

To describe variation among call types at the population level, we performed a principal components analysis (PCA). This returned three primary axes of variation (eigenvalues > 1) which together explained 67.8% of variation in acoustic properties (Table S2). Pulse duration, pulse interval, dominant frequency, and frequency of the 2^nd^ harmonic all loaded heavily on the first principal component (PC1; 37.8%), while call duration and number of pulses loaded heavily on the second principal component (PC2; 18.2%) (Table S2). The third principal component (PC3; 11.8%) contributed only subtly to variation among call types, with variation explained by call interval and pulse duration. Overlap between call types was evident across all individuals assayed (Fig. S2–3), suggesting that population-level patterns were not obscured by variation at the individual level.

To evaluate the predictive capacity of the PCs in assigning calls to their predefined social contexts, we next performed a discriminant function analysis (DFA) using PCs 1–3 as inputs. This approach correctly classified 72.2% of calls to their assigned social context. The chance-corrected classification success was significant (Weighted Cohen’s kappa = 0.566, *p* < 0.001). The strongest coefficients of the first linear discriminant were held by PC1 (the ‘frequency and pulse period’ factor), whereas PC2 (the ‘call length’ factor) contributed more to the second linear discriminant. The highest accuracy of classification according to this model was for advertisement calls (82.9%) followed by courtship calls (73.5%) and egg feeding calls (57.1%). Courtship calls were equally likely to be misclassified as advertisement (13.2%) or egg feeding calls (13.2%), and egg feeding calls were equally likely to be misclassified as advertisement (21.4%) or courtship calls (21.4%). While overall misclassification rates did not vary significantly across males (27.7 ±9.31%; χ^2^ = 21, *p* = 0.279), over half of all misclassified egg feeding calls (16/24) were attributed to three individuals: Ri.0181, Ri.0187, and Ri.0442 (Fig. S3). Given that egg feeding calls were sampled at the same grain (10 calls from a single bout of egg feeding) for each male, variation in parents or specific parenting conditions may explain variation in call distinctiveness.

## Discussion

Increased complexity of communication systems is predicted to co-evolve with the emergence of new forms of social interaction (1, 2, 6, 7), but this relationship has yet to be explored in the context of signals specialized for parental care. In this study, we characterized and compared calls of the biparental poison frog, *Ranitomeya imitator*, across social contexts in which calls either serve an ancestral communication function – advertisement or courtship – or a derived social function - parental care via coordinated, trophic egg feeding. We found that vocalizations produced in distinct social contexts differ in both acoustic properties and the amount of identity information contained in those properties. Crucially, calls elicited during egg feeding possessed unique spectral and temporal properties that were not observed in advertisement or courtship contexts and retained moderate identity information, suggesting that egg feeding calls were likely co-opted from existing elements but have since undergone evolutionary modification in form. Despite this, nearly half of all egg feeding calls were not sufficiently distinct to be distinguished from ancestral call types based on acoustic properties alone, implicating a role for multimodal communication in the coordination of parental behavior.

Studies of anuran acoustic communication recognize multiple common types of calls that function in distinct social contexts (15). Advertisement calls are by far the most studied call type because they serve multiple social functions (i.e., mate attraction, territory defense, and species delimitation) and are therefore taxonomically widespread (15). Virtually no attention has been paid to characterizing vocalizations deployed during parental care, in which communication between the sexes holds a special significance beyond simple mate attraction. We quantified nine acoustic properties of vocalizations elicited during egg feeding in *R. imitator* and compared these to the same properties quantified for advertisement and courtship calls. We found that egg feeding calls shared some properties with advertisement calls (i.e., longer than courtship calls with more pulses) and some with courtship calls (i.e., broader call bandwidths), supporting the prediction that egg-feeding calls evolved via co-option of elements from ancestral call types. This is not surprising, as male advertisement and courtship calls already function to elicit female responses. However, we also identified properties of egg feeding calls that were distinct from either ancestral call type (i.e., low dominant frequencies and especially short pulse durations coupled with long pulse intervals). Some of these differences in acoustic properties could be physiologically related to egg feeding calls being produced at lower amplitudes (i.e., “quieter”) because pairs are typically very close in space (<0.5 m) when males are leading females to tadpoles and stimulating them to feed (Movie S1). However, courtship calls are also “quiet” close-range signals, so the unique elements of egg feeding calls cannot be entirely explained as physiological by-products of low amplitude calls. Modification of temporal patterning is a common theme in the evolution of context-dependent signals (e.g., 22, 31) and we suggest the subtle differences observed in pulse durations and intervals reflect novel elements that are selectively maintained for the coordination of egg feeding.

A second major finding of our study was that the identity information conveyed in *R. imitator* vocalizations varies according to the social context in which they are deployed. By measuring calls repeatedly and across multiple social behaviors in the same males, we showed that advertisement calls contain the most individual identity information overall, with dominant frequency being the most reliable signature of male caller. Dominant frequency is relatively ‘static’ within males of many species (e.g., 11, 32) presumably because it is constrained by the morphology of the sound producing structures (35) and thus correlated with body size. Advertisement calls may be a signal or cue of individual identity used by rival males to recognize territory neighbors. In the case of mimic poison frogs, which are monogamous and pair bonding, individually distinctive advertisement calls could also be used by females to recognize their mate. If males benefit from being recognized by either territory neighbors or mates, selection could favor greater identity information in these calls to facilitate recognition. A complementary explanation is that courtship and egg feeding calls are typically produced over shorter distances than advertisement calls, such that other signal modalities (e.g., visual and/or chemosensory cues) may interact with auditory cues in mediating individual recognition. Indeed, the majority of acoustic properties examined were more dynamic (i.e., showed more within-individual variation) in courtship and egg feeding calls than in advertisement calls, which could reflect stronger selection for extreme values to enhance the motivational effects of these signals (33).

Finally, we evaluated whether subtle variation in the spectral range, temporal patterning, and individual distinctiveness was sufficient to quantitatively distinguish egg feeding calls from ancestral call types. In a discriminant function analysis, all three call types were classified above chance level (33%) from their acoustic properties alone; however, overlap between call types was considerable (Fig. 3A) and calls elicited during egg feeding were misclassified in nearly half of all instances. Thus, egg feeding calls may not be distinct enough to afford their own category within the vocal repertoire of *R. imitator* despite eliciting a novel, complex behavior. An intriguing observation can be made, however, in the patterns of misclassification. Egg feeding calls in our sample were equally likely to be misclassified as advertisement or courtship calls, suggesting that egg feeding calls borrow and recombine elements from multiple call types. Perhaps this complexity arises from the unique function of egg feeding calls, which may align selection on the response with both advertisement (i.e., must attract females to a specific location) and courtship calls (i.e., must motivate females to lay eggs). Additionally, females deciding to lay trophic eggs (which are typically laid in the water) versus eggs that will be fertilized (which are typically attached to a leaf above the water line or in a location away from a pool) are likely responding to their own physiological state and other contextual cues and signals as well as male signals. For one, tadpoles of this species vibrate vigorously against parental frogs to signal their hunger, which appears to be important in eliciting trophic eggs (22). Use of multimodal cues would also be consistent with acoustic studies of other frogs, which have shown that behavioral responsiveness to male calls is enhanced when females are also presented with visual (36) or contextual cues (37). Future studies should focus on female perception of male call variants in relation to female state and other contextual cues to better understand how selection on signalers and receivers could synergistically drive the co-option and modification of acoustic signals for trophic egg feeding.

**Figure 3.**
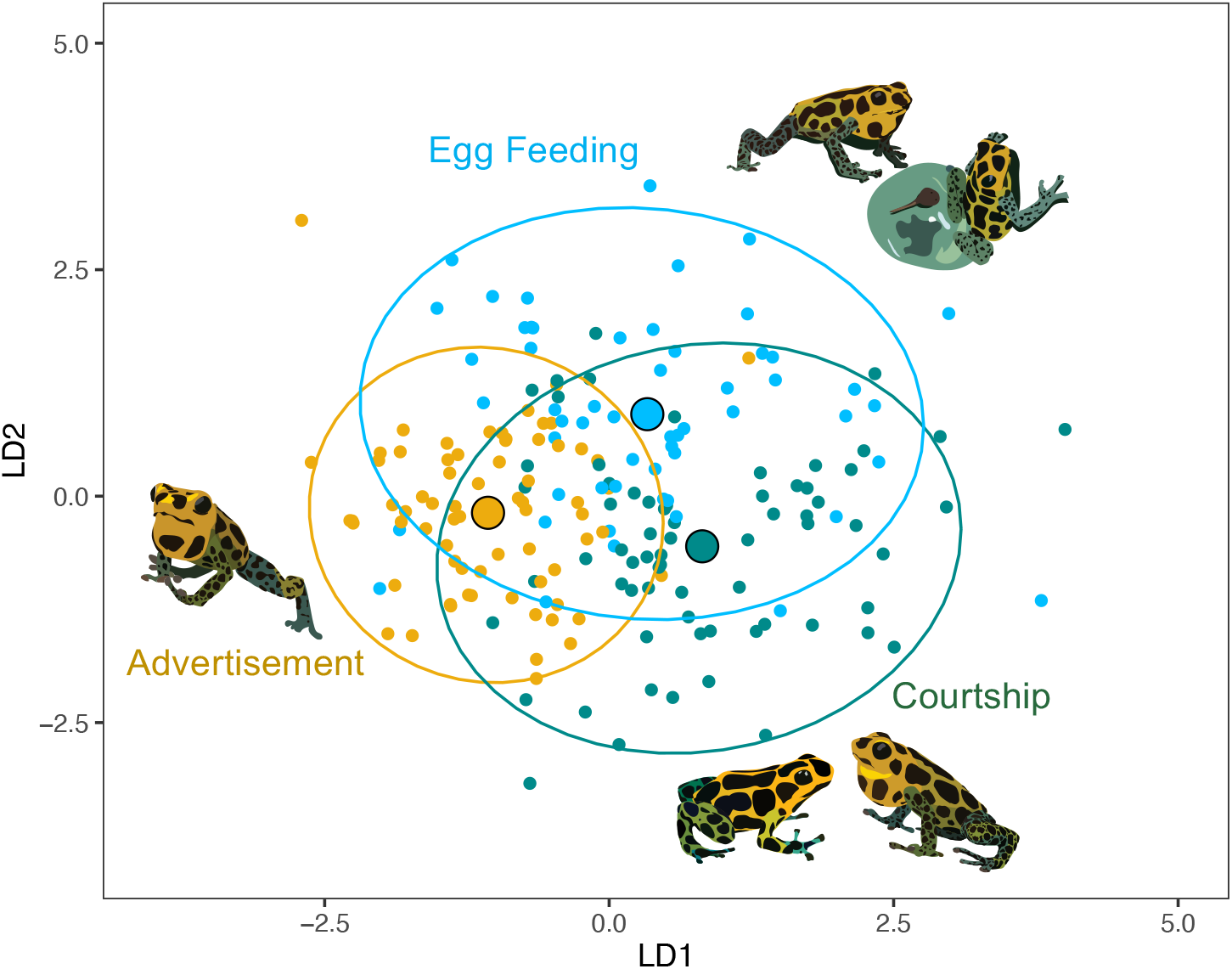
Classification success of vocalizations based on acoustic properties exceeds chance levels, despite considerable overlap. Plot of *Ranitomeya imitator* calls in two-dimensional space defined by the first two discriminant functions of a linear discriminant function analysis (DFA). Calls are clustered by call type (advertisement, courtship, and egg feeding) with centroids (large points) and 95% confidence intervals (ellipses) depicted for each type.

In conclusion, our study examined acoustic signals produced in a parental coordination context in poison frogs and contrasted these against ancestral signal types adapted to function in other facets of male-female communication. We found support for the prediction that extended functionality of the communication system in association with increased social complexity likely arose through co-option of pre-existing signals followed by evolutionary modification resulting in novel elements. Our results highlight the complexity of anuran communication systems and the need to characterize vocal repertoires across a greater diversity of social contexts. More generally, this study contributes to understanding how communication systems are evolutionarily fine-tuned to convey context-dependent information in increasingly complex social systems.

## Materials and Methods

### Animal Husbandry

A captive colony of *Ranitomeya imitator* descended from five localities in Peru (Veradero, Tarapoto, Huallaga, Sauce, Southern, and hybrids between these; Table S1) was maintained in temperature- (71.86 ± 2.79°C) and light- (12:12 h cycle) controlled rooms at the University of Illinois Urbana-Champaign (UIUC). Breeding pairs were housed in 12×12×18 inch glass terraria (©Exo Terra, Mansfield, MA, U.S.A) containing live plants, driftwood, sphagnum moss, leaf litter, and water-filled film cannisters for egg laying and tadpole deposition. Tank humidity (84.95 ± 3.08%) was maintained using a misting system (©Mist King, Ontario, Canada). Frogs were fed wingless *Drosophila* fruit flies dusted with vitamin supplements three times weekly. Only sexually mature individuals were used as focal subjects. All procedures were approved by the UIUC Animal Care and Use committee (Protocol #20147).

### Acoustic Recordings

Between July 2021 and May 2022, seven male *R. imitator* were recorded in their home terraria across the breeding cycle. A shotgun microphone (K6/ME66, ©Sennheiser, Wennebostel, Germany) was positioned above the screen covering the focal tank and connected to a handheld recorder (H4n Pro, Zoom, Tokyo, Japan); 44.1kHZ sampling rate, 16-bit resolution. For each recording, we documented the time of day and reproductive stage of the focal male by the following categories: (1) nonbreeding (pair did not currently have eggs or tadpoles); (2) brooding (frog had at least one egg but hatching had not yet occurred); or (3) tadpole (at least one tadpole was present in a deposition pool). Across all three stages, advertisement calls were distinguished from shorter courtship calls based on a published characterization of advertisement calls for this species (26). While advertisement calls produce a characteristic ‘trill’ sound and function in long-range mate attraction and territory defense, courtship calls are typically shorter and softer and function in close-range interactions with females (25, 26). To isolate vocalizations elicited in the unique social context of egg feeding and test whether these can be quantitatively distinguished from advertisement and courtship calls, we took video recordings (RLC-510A, ©Reolink, New Castle, DE, U.S.A.) concurrently with audio recordings during the tadpole stage. Male *R. imitator* stimulate female partners to lay trophic eggs by leading them to tadpole deposition pools and maintaining close and/or tactile contact while calling continually (21). Thus, egg feeding calls were identified as calls coinciding with males moving towards tadpole deposition pools, leading females to pools, sitting on or near the pools, and culminating with the arrival of females at pools (e.g., Movie S1). Calls recorded during the tadpole stage while males were away from pools and/or oriented towards egg laying cannisters were identified as either advertisement or courtship calls, based on the same criteria used during nonbreeding and brooding stages.

### Call Filtering and Analysis

To examine variation within and among the three call types, ten calls of each type for each individual were selected from available recordings (Dataset S1). Very low-amplitude or low-quality calls, or calls that could not be clearly classified between the three types on the basis of social context or previous published accounts were excluded. Raw recordings of calls were trimmed in Audacity v. 3.0.2 (Audacity Team, 2021). Analysis of calls was performed in Raven Pro v. 1.6.1 (Cornell Lab of Ornithology, 2022). In total, we measured nine acoustic properties of each call. We measured six temporal properties using the entire call or part of the call (i.e., a single pulse or interpulse interval) as the unit of measurement in the waveform view in Raven. These included call duration (time in seconds from the first pulse to the last note; Fig. S1A), inter-call interval (time in seconds to the next call within the same bout, with bout defined as calls occurring within one minute of each other; Fig. S1A), number of pulses, pulse duration (time in seconds from the beginning of a note to the end; Fig. S1B), pulse interval (time in seconds from the end of one note to the start of the next note), and pulse rate (the reciprocal of pulse period, or the full time from the beginning of one note to the beginning of next consecutive note; Fig. S1B). Pulse duration, pulse interval, and pulse rate were calculated by taking the average of three notes within the call, excluding introductory notes. Three spectral properties were also quantified using the entire call as the unit of measurement, including the dominant frequency of the call (in Hz; Fig. S1C) and its bandwidth (90%; Fig. S1C). In most cases the dominant frequency corresponded to the second harmonic (5-6 kHz) but occasionally the fundamental frequency (<3 kHz) was dominant and in a few cases, a broad frequency peak in the range of 3-4 kHz was observed and the harmonic structure was unclear (Fig. 1; Fig. S1). Thus, the frequency of the second harmonic was also quantified for each call.

### Statistical Analyses

All statistical analyses were performed in R Studio v. 1.1.318 (R Core Development Team, 2020). We first summarized each of nine acoustic properties by their assigned call type. We tested for differences between call types using Type III ANOVAs followed by post-hoc testing. Model testing was carried out using the packages ‘lmerTest’ (38) and ‘emmeans’ (39), with reproductive stage specified as a covariate and individual ID specified as a random effect in all models. For metrics that were collected in multiples per call (i.e., pulse duration, pulse interval, and pulse rate), we also specified call ID nested within individual ID as a random effect. Some variables (inter-call interval, pulse interval, pulse rate, and all spectral properties) were log-transformed prior to analysis to better meet assumptions of normality and homogeneity of variance, which we evaluated with analyses of random effects and residuals.

We next evaluated within- and among-individual variation for each call type for each of the nine acoustic properties. For each male, we calculated a within-individual coefficient of variation (CV_w_ = individual SD/individual mean × 100) and report the mean CVw across males. We computed the among-individual coefficient of variation (CV_a_ = grand SD/grand mean × 100) and report the ratio of among-individual variability (CV_a_) to within-individual variability (CV_w_) as the potential for individual coding (PIC; 43). Values of PIC greater than one indicate that a call property varies more among individuals than within individuals. To further describe individual distinctiveness of calls, we calculated Beecher’s Information Statistic (*H_s_;* 44), which expresses the overall amount of individual identity information contained in calls. To control redundancy due to correlations among measurements, data were transformed into uncorrelated variables using principal components analysis (PCA) and all principal components were retained to calculate *H_s_* (30). PCAs were implemented with the ‘prcomp’ function in R after transforming pooled variables (pulse duration, pulse interval, and pulse rate) into averages to ensure equality of sample sizes and scaling all input variables.

Finally, to describe variation among call types at the population-level and evaluate the success rate of classifying calls to the correct call type (advertisement, courtship, or egg feeding), we took a two-step multivariate approach. First, we performed PCA as described above. Input variables were prepared by converting acoustic properties into z-scores scaled to the mean of the individual, which controlled for variation among males. Second, principal components with eigenvalues > 1 were extracted as inputs into a discriminant function analysis (DFA). We implemented this test using the linear discriminant analysis (‘lda’ function) in the package ‘MASS’ (41). Chance-corrected classification success (Cohen’s kappa) was calculated following the approach of (42) in the R package ‘vcd’ (43). Differences in classification success among males were statistically evaluated using a Chi-Squared test of independence.

## Acknowledgments

We thank Mark Bee for helpful advice on bioacoustics equipment and approaches in early project design stages. We thank the Fischer Lab for constructive feedback on earlier drafts of our manuscript. This study was supported by a National Science Foundation Postdoctoral Research Fellowship in Biology to J.B.M. (#2010649), at University of Illinois Research Board Grant to E.K.F. (RB21025), and University of Illinois Urbana-Champaign laboratory start-up funds to E.K.F.

