## Supplementary material for "Co-option of ancestral signal elements in the evolution of a cooperative parental behavior": Table S1, Table S2, Fig. S. 1, Fig. S1A, Fig. S1B, Fig. S1C, Fig. S2

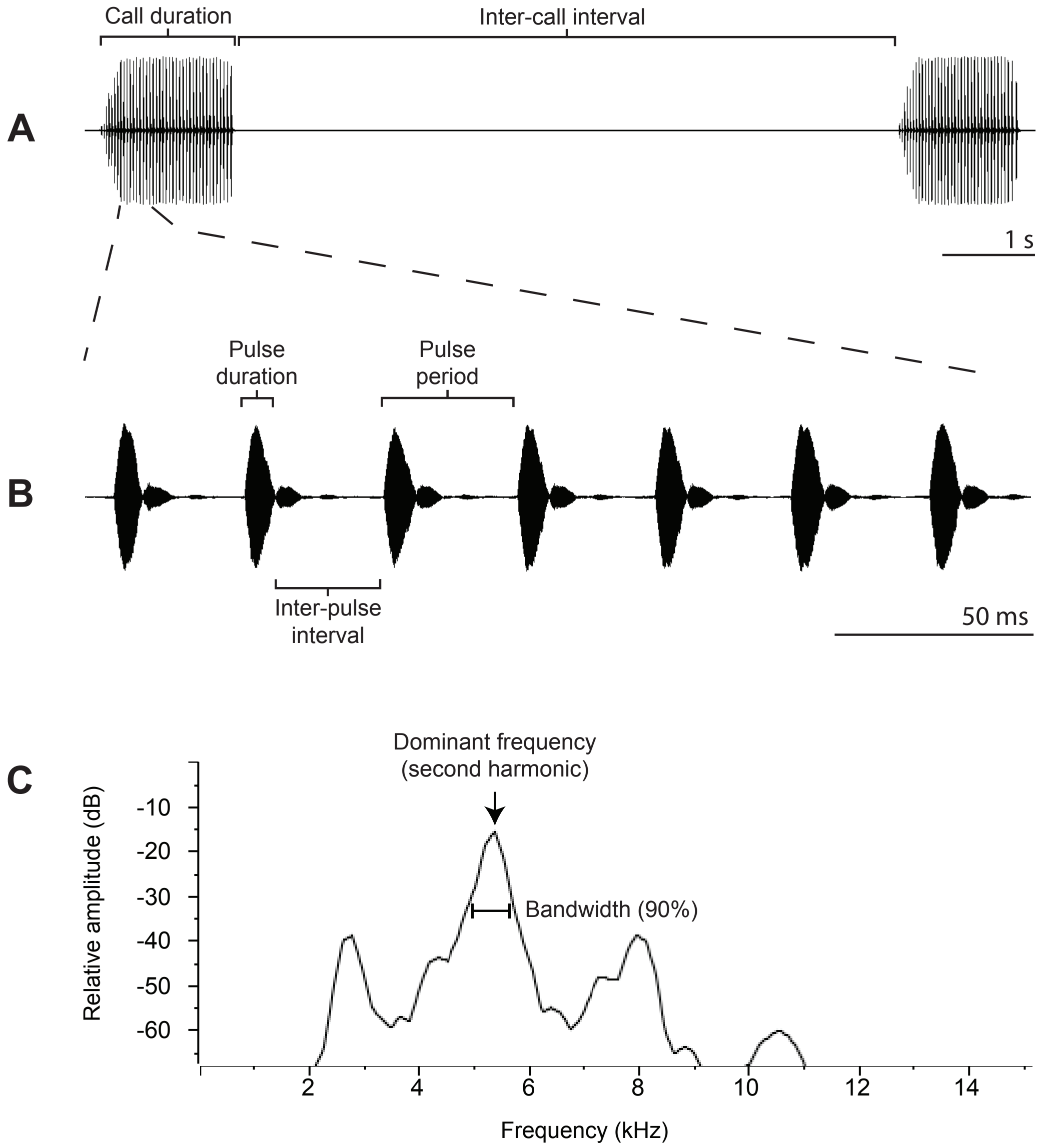


**Fig. S1.** Illustration of the acoustic properties of calls that were analyzed for this study. (A) Temporal properties were measured as call duration (time from the onset of the first pulse in a call to the offset of the last pulse in a call) and inter-call interval (time from the offset of the last pulse in a call to the onset of the first pulse in a subsequent call). (B) Temporal properties of pulses within a call were measured as pulse duration (time from onset to offset of an individual pulse) inter-pulse interval (time from offset of a pulse to onset of the subsequent pulse), and pulse period which was the sum of pulse duration and pulse interval. (C) Spectral properties were measured as dominant frequency (frequency peak with the highest amplitude, a.k.a. “max frequency” in Raven) and 90% frequency bandwidth (difference in frequency between the 5% and 95% frequencies, representing 90% of the sound energy, a.k.a. “bandwidth 90%” in Raven). In this example call, the dominant frequency is the $2^{nd}$ harmonic, but in cases where the fundamental frequency was dominant, we also measured the frequency of the $2^{nd}$ harmonic. All measurements were taken in Raven Pro v. 1.6.1.

.

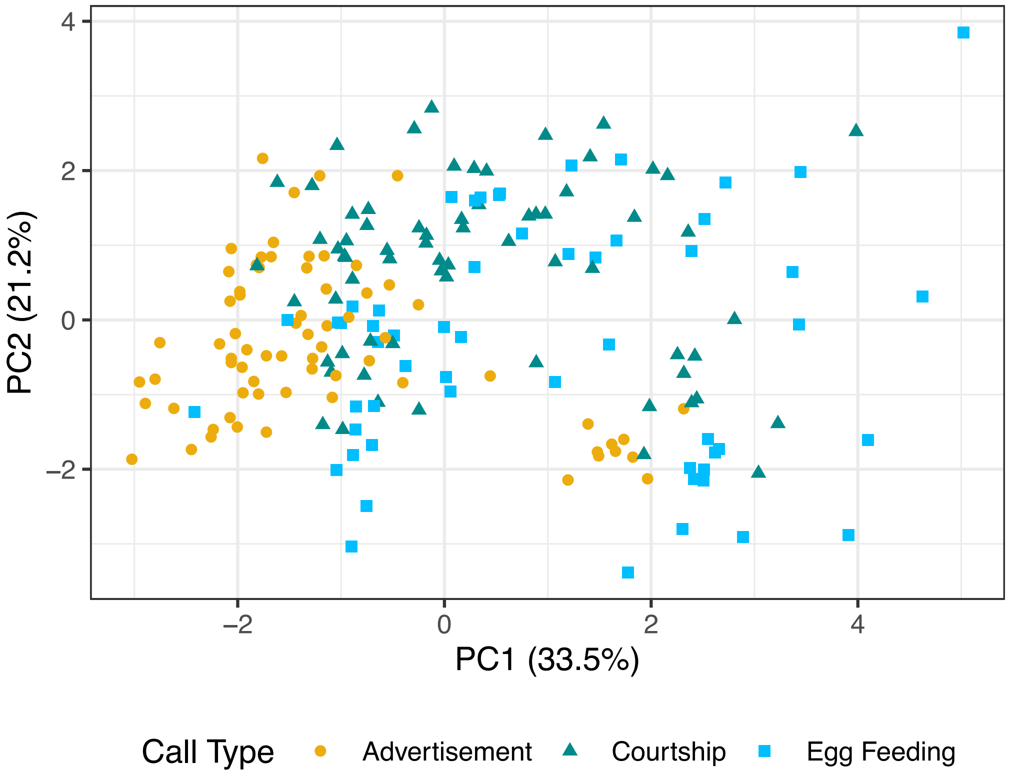


**Fig. S2.** Plot of 196 calls in two-dimensional space defined by the first two axes of a principal components analysis (PCA). The PCA was performed with acoustic properties standardized to the population mean to compare individual distributions. Colors represent the three types of calls – advertisement (yellow), courtship (green), and egg feeding (blue).

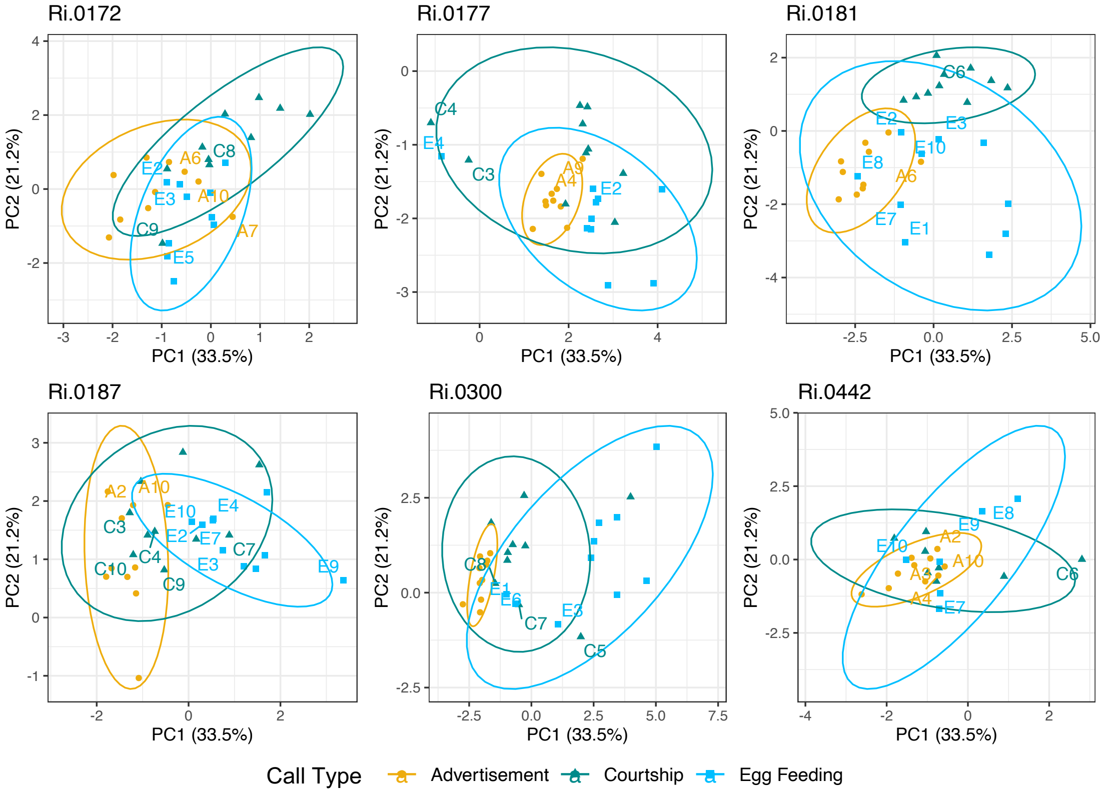


**Fig. S3:** Plot of calls in two-dimensional space defined by the first two axes of a principal components analysis (PCA). Each panel depicts calls of an individual male with the exclusion of Ri.0188, for which egg feeding calls could not be obtained. Ellipses represent 95% confident intervals for each of the three types of calls – advertisement, courtship, and egg feeding. Calls that were misclassified in a discriminant function analysis are labeled for each male.

**Table S1. Localities of origin of seven focal males used in this study.**

| Male ID | Locality of Origin |
|---|---|
| Ri.0172 | Unknown |
| Ri.0177 | Varadero x Tarapoto |
| Ri.0181 | Varadero x Tarapoto |
| Ri.0187 | Varadero x Tarapoto |
| Ri.0188 | Sauce x Huallaga |
| Ri.0300 | Southern |
| Ri.0442 | Varadero |

**Table S2.** Summary of three principal components (PCs) with eigenvalues > 1 resulting from principal components analysis of 196 calls spanning seven males and three call types. Variables with the largest contributions to each PC are shown in bold.

| **Variable** | **PC1 (37.8%)** | **PC2 (18.2%)** | **PC3 (11.8%)** |
|---|---|---|---|
| Call Duration (s) | -0.264 | **-0.610** | 0.218 |
| Interval to next call (s) | -0.137 | -0.281 | **-0.553** |
| No. of Pulses | -0.356 | **-0.482** | 0.199 |
| Pulse Duration (s) | **-0.400** | 0.085 | **-0.464** |
| Pulse Interval (s) | **0.452** | -0.239 | 0.343 |
| Pulse Rate (notes/s) | -0.297 | 0.384 | 0.112 |
| Dominant Frequency (Hz) | **-0.372** | 0.227 | 0.345 |
| Frequency of $2^{nd}$ Harmonic (Hz) | **-0.366** | 0.180 | 0.375 |
| Bandwidth 90% (Hz) | 0.245 | 0.144 | 0.043 |
